# Discordant attributes of structural and functional connectivity in a two-layer multiplex network

**DOI:** 10.1101/273136

**Authors:** Sol Lim, Filippo Radicchi, Martijn P van den Heuvel, Olaf Sporns

## Abstract

Several studies have suggested that functional connectivity (FC) is constrained by the underlying structural connectivity (SC) and mutually correlated. However, not many studies have focused on differences in the network organization of SC and FC, and on how these differences may inform us about their mutual interaction. To explore this issue, we adopt a multi-layer framework, with SC and FC, constructed using Magnetic Resonance Imaging (MRI) data from the Human Connectome Project, forming a two-layer multiplex network. In particular, we examine whether node strength assortativity within and between the SC and FC layer may confer increased robustness against structural failure. We find that, in general, SC is organized assortatively, indicating brain regions are on average connected to other brain regions with similar node strengths. On the other hand, FC shows disassortative mixing. This discrepancy is apparent also among individual resting-state networks within SC and FC. In addition, these patterns show lateralization, with disassortative mixing within FC subnetworks mainly driven from the left hemisphere. We discuss our findings in the context of robustness to structural failure, and we suggest that discordant and lateralized patterns of associativity in SC and FC may explain laterality of some neurological dysfunctions and recovery.

## 1 Introduction

The relationship between structural connectivity and functional connectivity has attracted much attention in recent years (Park & Friston, 2013; Sporns, 2013b; Damoiseaux, 2017; Uddin et al., 2011; Mišić *et al*., 2016). Yet despite numerous empirical (Uddin *et al*., 2011; Betzel *et al*., 2014) and computational studies (Sporns, 2013b; Park & Friston, 2013) the nature of their interaction remains only incompletely understood. Several studies have suggested that functional brain networks are constrained by the underlying structural connectivity (Sporns, 2013a), and brain-wide comparisons have supported the idea that functional connectivity (FC), measured in the resting state, and structural connectivity (SC) are in general statistically correlated (Skudlarski *et al*., 2008). For example, when there is a strong anatomical connection between two areas of the brain, the corresponding functional connection is likely to be strong as well, but the inverse is not always the case (Koch *et al*., 2002; Damoiseaux & Greicius, 2009; Messé *et al*., 2014; Mišić *et al*., 2016; de Pasquale *et al*., 2017; Honey *et al*., 2010; Honey *et al*., 2009). A number of computational models have successfully reproduced some features of empirical functional connectivity, including models based on large-scale dynamics (Breakspear, 2017; Deco *et al*., 2013) or graph theory metrics (Goñi *et al*., 2014). While most studies have emphasized the statistical association between SC and FC, important differences and discrepancies remain (Honey *et al*., 2009; Messé *et al*., 2014). This may be expected given that fMRI and dMRI measure different signals and use different statistical approaches for estimating pairwise connections between regions of interest (ROIs). Whereas SC estimates a direct relationship or path between two brain regions, measurements of FC, for instance, estimated by Pearson’s moment correlation coefficients, incorporate both direct and indirect relationships between two nodes influenced by other brain areas (van den Heuvel & Sporns, 2013; Zalesky *et al*., 2012).

However, not many studies have focused on fundamental topological differences in the network organization of SC and FC, and on how these differences may provide insight into their mutual interaction. To explore this issue, we adopt a multi-layer framework, with SC and FC forming a multiplex network (Buldyrev *et al*., 2010; Boccaletti *et al*., 2014; Kivelä *et al*., 2014). What are the fundamental topological differences that underpin FC and SC, and are these differences biologically meaningful? Are there potential benefits that might arise from topological differences among these two different types of brain networks? As previous studies have shown, SC and FC are intricately (and non-trivially) linked – for example, by demonstrating that FC arises from underlying anatomical connections. Thus, if any systematic/consistent topological differences exist between these two networks, are they mere by-products of the generative process or could they represent biologically meaningful features of multi-layer organization that carry benefit or enhance overall functionality?

Specifically, we will examine the possibility that node strength assortativity between the SC and FC layers may confer increased robustness to the multiplex system. Degree assortativity has been extensively studied in the context of network robustness (Newman, 2003; Callaway *et al*., 2000; Noldus & Van Mieghem, 2015). In isolated networks, assortativity stands for correlation among nodes features (e.g., degree) of directly connected nodes (Newman, 2003). A network is said to be assortative if its connectivity pattern is such that high-degree nodes are frequently attached to other high-degree nodes, and low-degree nodes are preferentially connected to other low-degree nodes. Assortative networks are generally resilient against the random removal of nodes and edges (Newman, 2003; Pechenick *et al*., 2012; Vázquez & Moreno, 2003). In multiplex networks, correlations among nodes features can be measured both within- and between-layers (Nicosia & Latora, 2015; de Arruda *et al*., 2016). The two types of correlations provide different information about the robustness of the interdependent system. In the absence of any correlations between layers, it is well known that an interdependent network undergoes a sudden percolation transition (Buldyrev *et al*., 2010). An increased within-layer degree assortativity decreases the robustness of the network in terms of the percolation threshold (Zhou *et al*., 2012). On the other hand, positive values of between-layer correlations generally mitigate the abrupt nature of the transition, making the system more robust. Examples include degree-degree correlations (Reis *et al*., 2014), edge overlap (Cellai *et al*., 2013; Min *et al*., 2015; Radicchi, 2015; Radicchi & Bianconi, 2017; Baxter *et al*., 2016), clustering and spatial coordinates (Danziger *et al*., 2016; Kleineberg *et al*., 2016; Kleineberg *et al*., 2017).

Robustness is an important feature of brain networks (Petrosini, 2017; Aerts *et al*., 2016; Bullmore & Sporns, 2012). In many cases, FC network patterns appear to maintain large-scale patterns and functionality even in the face of serious disruptions or disturbance of underlying SC. Can a multi-layer model shed light on the network basis for these observations? Here, we investigate if these findings from theoretical investigations and non-biological networks carry over to human brain networks derived from Magnetic Resonance Imaging (MRI) data from the Human Connectome Project (Van Essen *et al*., 2013). SC was constructed based on diffusion MRI and tractography and FC was estimated using regularized partial correlation coefficients with the elastic net. The two layers (SC and FC) are coupled by creating links on pairs of corresponding nodes in the two layers, thus creating a multiplex network. We examine assortative mixing by strength both within SC and FC and between the two layers. We divided FC and SC into 7 subnetworks according to a canonical resting-state partition (Yeo *et al*., 2011). We find that coupled structural and functional human brain networks exhibit a combination of similarities and differences. In addition, we find heterogeneous strength-strength correlations across the two layers and within FC and SC subnetworks, as well as between the left and right hemispheres. Our findings may offer clues to understand why some brain networks are more vulnerable to or more resilient against functional disruption due to brain disorders or injuries.

## 2 Methods and materials

### 2.1 Data and data processing

The dataset was provided by the Human Connectome Project (HCP; http://www.humanconnectome.org) from the Washington University-University of Minnesota (WUMinn) consortium (Van Essen *et al*., 2013), acquired using a modified 3T Siemens Skyra scanner with a 32-channel head coil. Resting-state fMRI data in an eyes-open condition were collected for approximately 14 min (1,200 time points) with TR = 720 *ms*, TE = 33.1 *ms*, flip angle = 52°, voxel size = 2 *mm* isotropic, and FOV = 208 × 180 *mm*^2^ and 72 slices. The data were acquired with opposing phase encoding directions, left-to-right (LR) in one run and right-to-left (RL) in the other run. Scanning parameters of a *T*_1_-weighted structural image were TR = 2,400 *ms*, TE = 2.14 *ms*, flip angle = 8°, voxel size = 0.7 *mm* isotropic, FOV = 224 × 224 *mm*^2^ and 320 slices. Diffusion-weighted images (DWI) were acquired with 270 gradient directions with *b*-values 1000, 2000, 3000 *s/mm*^2^, two repeats, and in a total of 36 *b*_0_ scans: TR = 5520 *ms*, TE = 89.5 *ms*, flip angle = 78°, FOV = 210 × 180 *mm*^2^, 111 slices, and voxel size = 1.25 *mm* isotropic. A T_1_-weighted structural image was acquired with TR = 2400 *ms*, TE = 2.14 *ms*, flip angle = 8°, FOV = 224 × 224 *mm*^2^, 320 slices, and voxel size = 0.7 *mm* isotropic. From the minimally preprocessed DWI data, white matter fibers were reconstructed using generalized q-sampling imaging (Yeh *et al*., 2010) and deterministic streamline tractography (de Reus van den Heuvel, 2013; de Reus & van den Heuvel, 2014; van den Heuvel *et al*., 2015; van den Heuvel *et al*., 2016). Our study included 484 participants in total from the Q4 release of HCP data.

### 2.2 Structural connectivity (SC) and functional connectivity (FC)

Both structural networks and functional networks consisted of 219 cortical nodes using a subdivision parcellation (Cammoun *et al*., 2012) of the Desikan-Killiany atlas (Desikan *et al*., 2006). For the structural networks, the edge weights were defined by the streamline count between two ROIs derived from diffusion MRI tractography and the edge weights for the functional networks were estimated as regularized partial correlation coefficients (see Section 2.6 for details).

### 2.3 Interdependent relationship between SC and FC

We model the interdependency of SC and FC using a multi-layer network approach (Boccaletti, 2004; Kivelä *et al*., 2014). SC and FC form two separate layers that are linked by multiplex coupling, such that a node (ROI) in one layer is connected to the same node in the other layer in a one-to-one correspondence (Figure 1, left). Building on previous work that has shown significant interactions between SC and FC (Park & Friston, 2013; Sporns, 2013b; Damoiseaux, 2017; Uddin *et al*., 2011; Mišić *et al*., 2016), the multi-layer approach is designed to consider SC and FC as interdependent networks with a one-to-one correspondence. In this study, we consider a two-layer multiplex network with unweighted dependency links between layers for simplicity with a minimal set of assumptions; however, one could extend the model to represent a more general multi-layer network framework for future studies.

**Figure 1:**
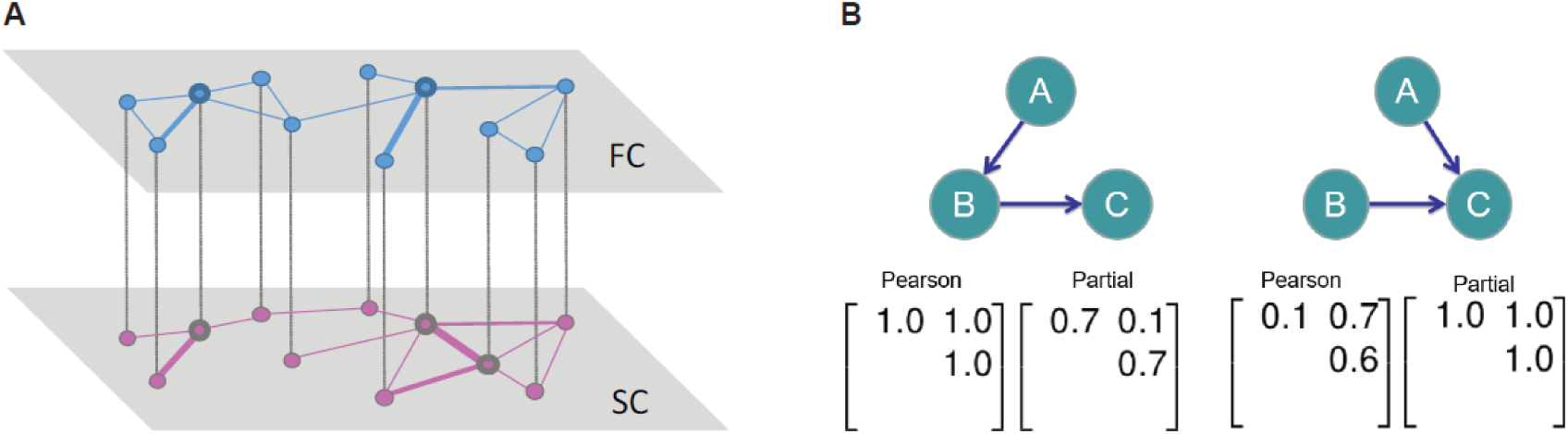
(A) A schematic representation of a brain multiplex network, where the networks of functional connectivity (FC) and structural connectivity (SC) are coupled via one-to-one match between corresponding region of interests; hub nodes in the SC (denoted as bigger circles) may not coincide with high-degree nodes in the FC; nodes linked in SC may not coactive strongly, resulting in the absence of the edge in FC. (B) Limitations of Pearson’s correlation and partial correlation. Depending on the underlying neuronal circuits both approaches can result in undesirable connection weights in the functional graph. (Left) A case where Pearson’s correlation coefficient fails to disregard a non-existent connection between node A and C. The visualization stands for the underlying probabilistic graphical model among the variables A, B, and C, with connections standing for dependencies among pairs of variables. The two matrices contain the coefficients of Pearson’s correlation and partial correlation coefficients, respectively. Rows and columns of the matrices refer to alphabetically ordered pairs of variable. Given their symmetry, we show only the upper-triangle of the matrices. (Right) A case where the partial correlation coefficient counter-intuitively imposes a high weight on the connection between node A and B due to the dependence on C. (Adapted from Nie 2015)

### 2.4 Estimation of functional connectivity from resting-state fMRI

Many different measures have been used to estimate or construct functional brain networks (Bullmore & Sporns, 2009; Deco *et al*., 2011; Friston *et al*., 2003). Among them, the Pearson’s moment correlation coefficient has been the most popular choice among brain researchers, and, despite its simplicity, it has provided valuable information regarding the intrinsic functional organization of the brain (Wang *et al*., 2014). However, other measures for estimating pairwise functional connectivity do exist, and they probe different aspects of dynamic interactions. Alternative choices include partial correlation coefficients, coherence measures that estimate linear relationships considering direct/indirect coupling effects in either the time domain or in the frequency domain, or non-linear measures such as mutual information, second-order maximum entropy or generalized synchronization (Simpson *et al*., 2013; Wang *et al*., 2014). In addition to the aforementioned seed-based definition of ROIs for functional networks, spatial ICA-based functional networks are also widely used (van de Ven *et al*., 2004; Calhoun *et al*., 2001; Smith *et al*., 2013). Importantly, network properties of FC may differ depending on which dependency/synchrony measure one chooses as each measure captures different aspects of the functional network (Wang *et al*., 2014). Spatial ICA and seed-based FC have shown to be similar in certain cases (Van Dijk *et al*., 2010) and both have advantages and disadvantages (for more quantitative comparison between the two methods, see (Joel *et al*., 2011)). Importantly, network properties of FC may differ depending on which dependency/synchrony measure or which type of derived FC one chooses to construct as each method captures different aspects of the functional network (Simpson *et al*., 2013; Wang *et al*., 2014; Zalesky *et al*., 2012; Liang *et al*., 2012; Jalili, 2016). Here, we use seed-based FC in order to make a direct connection between SC and FC, in particular, we use regularized partial correlation coefficients based on a regression approach (Krämer *et al*., 2009). In the following section, we elaborate on why we chose to use regularized partial correlation coefficients.

### 2.5 Full correlation versus partial correlation

The utility of a partial correlation approach to functional brain networks derives from the capacity to remove indirect effects due to remote linear effects propagated from other regions (van den Heuvel *et al*., 2008; Supekar *et al*., 2010; Ryali *et al*., 2012; Hampson *et al*., 2002; Marrelec *et al*., 2006; Smith *et al*., 2011). However, using partial correlation coefficients entails some other issues, such as requiring number of observations larger than the number of ROIs, potential overfitting and less stable estimation (Ryali *et al*., 2012; Smith, 2012; Krämer *et al*., 2009; Nie *et al*., 2015; Friedman *et al*., 2008; Huang *et al*., 2010; Lee *et al*., 2011; Peng *et al*., 2009; Meinshausen & Bühlmann, 2010; Varoquaux *et al*., 2010). On the positive side, partial correlation coefficients could estimate connection strengths between two brain regions that are conditionally independent with a small coefficient, reducing or removing indirect connections (often referred to as spurious connections) (Figure 1, left). Nonetheless, when the underlying structure happens to involve conditional dependence, in other words, ‘explaining away’ phenomenon in the Bayesian modelling literature (probabilistic graphical models) (Pearl, 1988), estimating partial correlation coefficients can cause Berkson’s paradox, inducing a ‘spurious’ connection, which will not happen when we use full correlation coefficient estimation (Figure 1B, right) (Berkson, 1946). This can be partially solved by using regularized partial correlation coefficients (Nie *et al*., 2015).

### 2.6 Regularized partial correlation coefficients using elastic net

Regularized partial correlation coefficient estimation has been proposed for constructing functional networks based on resting-state fMRI to overcome the limitation of partial correlation coefficient estimation, while measuring direct relationships between two brain areas(Ryali *et al*., 2012; Smith, 2012; Krämer *et al*., 2009; Nie *et al*., 2015; Friedman *et al*., 2008; Huang *et al*., 2010; Lee *et al*., 2011; Peng *et al*., 2009; Meinshausen & Bühlmann, 2010; Varoquaux *et al*., 2010) and, more commonly, for constructing gene association networks (Krämer *et al*., 2009). Estimation of regularized partial correlation coefficients has been carried out by applying regularization on Gaussian Graphical Models (GGMs), which can be represented as a graph with edges estimating conditional dependence between nodes (Whittaker, 2009; Krämer *et al*., 2009). Of those regularized models, the elastic net has been shown to be a good model to estimate resting-state functional connectivity (Ryali *et al*., 2012). Although providing sparser solutions which do not require further statistical thresholding, *L*_*1*_-norm regularization can only identify a number of functional connections that is less than or equal to the number of observations (time points) and can detect only a subset of connections when the time series are highly correlated (Zou & Hastie, 2005). On the other hand, *L*_2_-norm regularization does not shrink small values of coefficients to zero, and hence we may not achieve the desirable level of sparseness of the network (Zou & Hastie, 2005). We can overcome these limitations by using the elastic net regression, which uses penalization of both *L*_*1*_ and *L*_*2*_ norms, or a linear combination of *L*_*1*_ and *L*_*2*_ norm regularization by solving the following problem (Friedman *et al*., 2010).

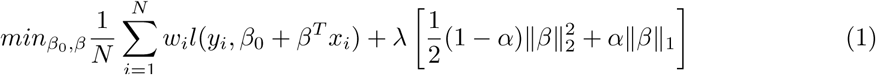

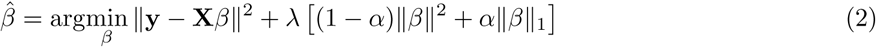

While trying to minimize our objective function (eq.1), we need to optimize our parameters *λ* and *α*. *λ* controls the overall penalization of the model and *α* determines how much we would put weight on *L*_1_-regularization compared to *L*_2_ regularization. For example, if *α* is 1, our model becomes LASSO or *L*_1_ regularization model, which will give us the sparsest graph. There are several methods to identify optimal parameter values such as grid search, which is slow and unstable because grid density affects the accuracy and depends on heuristic choices for parameter ranges. Alternatively, one could also use a stability selection method, which aims to control the false discovery rate (Meinshausen & Bühlmann, 2010) to determine the proper amount of regularization. Here, we made use of the interval search EPSGO algorithm to tune our parameters *λ* and *α* based on 10-fold cross-validation. This algorithm learns a Gaussian process model of the loss function surface in parameter space and samples at points where the expected improvement criterion is maximal (Sill *et al*., 2014; Frohlich & Zell, 2005; Jones *et al*., 1998). After calculating regularized *β*s (coefficients of predictors) with the optimized regularization parameters, we obtain partial correlation coefficients from *β*s (Whittaker, 2009; Krämer *et al*., 2009).

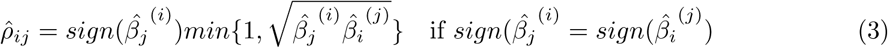

otherwise 0
where 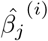 is the regularized estimate of *β*s between brain region *i* and the rest of the brain regions except the region *i*, which ensures the partial correlation coefficients are well-defined and in the interval [-1, 1].

### 2.7 Assortativity mixing within and between layers

Assortativity quantifies the tendency for nodes to connect to other nodes that are similar in some way (Newman, 2003; Noldus & Van Mieghem, 2015). For example, modularity Q (Newman, 2003; Newman & Girvan, 2004) is an assortativity-based measure which expresses the actual connection density of nodes within the same community compared to the value of the connection density expected in a suitably defined null model. We can also measure the tendency of ‘similar’ nodes being connected to each other (actual vs. expected) based on some scalar nodal attribute such as degree, or betweenness. One of the most common cases where we define assortative mixing according to a scalar quantity is assortativity mixing by degree; positive degree assortativity implies that high-degree nodes are preferentially connected to high-degree nodes on average and low-degree nodes mainly connect to low-degree nodes on average. Since FC edges can carry either positive or negative weights, we considered a version of assortativity that takes into account node strengths, called node strength assortativity. The strength of a node is defined by the sum of its all weights (Barrat *et al*., 2004). In FC a node’s strength is close to zero when its neighbors maintain positive and negative weights that nearly balance out. In this study, high values of the assortativity for the FC layer reflect a connectivity pattern where nodes with high strength are tendentially connected, through positively valued edges, to nodes with high strength. We did not weight the connection strength between two nodes, rather we measured the strength correlation between two nodes. In other words, we calculated Pearson’s correlation coefficient between a pair of nodes based on their strengths:

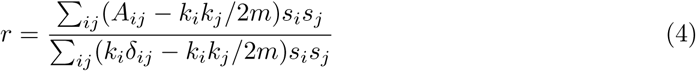

Where *k*_i_ is the degree of node *i* and 2*m* = ∑_*i*_ *k*_*i*_ *A_ij_* = 1 if a connection between nodes *i* and *j* exists, otherwise *A_ij_* = 0. *s*_*i*_ = ∑_*i*_ *k*_*i*_ *w*_*ij*_ is the sum of the weights of all connections departing from node *i*. *δ_ij_* = 1 if *i* = *j* and *δ_ij_* = 0, otherwise. The numerator of Eq. 4, is the covariance of the pair of *s*_*i*_ and *s*_*j*_ on the edge (*i*, averaged or all pairs of edges. We define the mean *µ* of the *s*_*i*_ at the end of an edge as 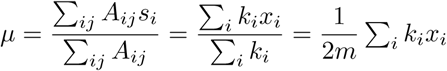, which is is the average over edges rather than over all vertices. Then the covariance of *s*_*i*_ and *s*_*j*_ over edges is the following.

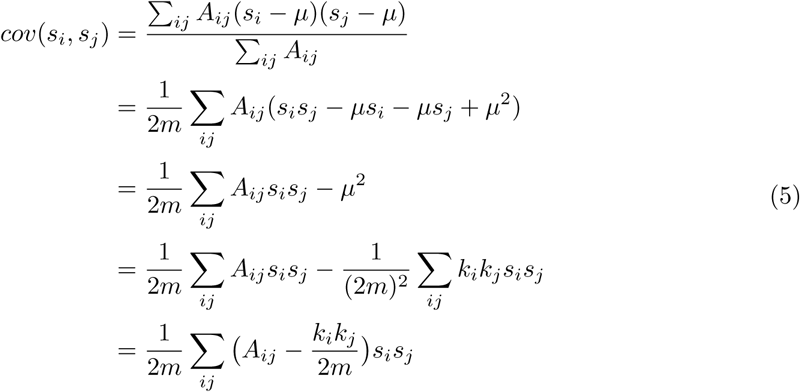

To normalize this, we devide Eg.5 by the following equation where all edges connect two nodes with the equal values of *s*_*i*_. When we replace *s*_*j*_ with *s*_*i*_, we have

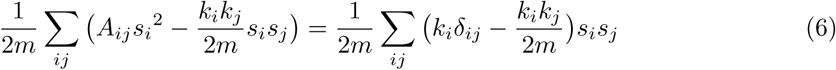

Thus, our node strength assortativity is *r* = *cov*(*s*_*i*_, *s*_*j*_)/*var*(*s*_*i*_), which is Eq.4 (Newman, 2010). We measure the coefficient *r* within the SC and FC layers. Furthermore, we subdivide our networks into Yeo’s 7 networks (Yeo *et al*., 2011) and investigated strength assortativity within each subnetwork for FC and SC as well as the strength assortativity between FC and SC for each network. In addition, we compared the left and the right hemispheres for each subnetwork.

### 2.8 Statistical analysis

Paired permutation test was used to compare the left and right hemisphere median differences for each subnetwork (Strasser & Weber, 1999) by approximating the exact conditional distribution using conditional Monte Carlo procedures (10000 permutations) and corrected by Bonferroni method (Dunn, 1959). A two-tailed test was used with alpha level 0.05 and adjusted p-values were reported based on Bonferroni correction. All statistical tests and calculations were performed either in Matlab R2016b (Mathworks Inc., Natick, MA) and R with R packages (http://www.R-project.org/) (R Core Team, 2013).

## 3 Results

### 3.1 Assortative within-layer structural connectivity (SC), disassortative within-layer functional connectivity (FC) and between-layer assortativity (SC and FC)

We first calculated node strength assortativity in each individual data set across the entire cerebral cortex. Within each layer, FC was characterized by disassortative connectivity (i.e., negative assortativity, Figure 2C Left), while (SC showed assortative connectivity with a somewhat broader variability among individuals (Figure 2C Right). The coupling between FC and SC assuming multiplexity between the two layers demonstrated a weak but positive assortativity in general (Figure 2D).

**Figure 2:**
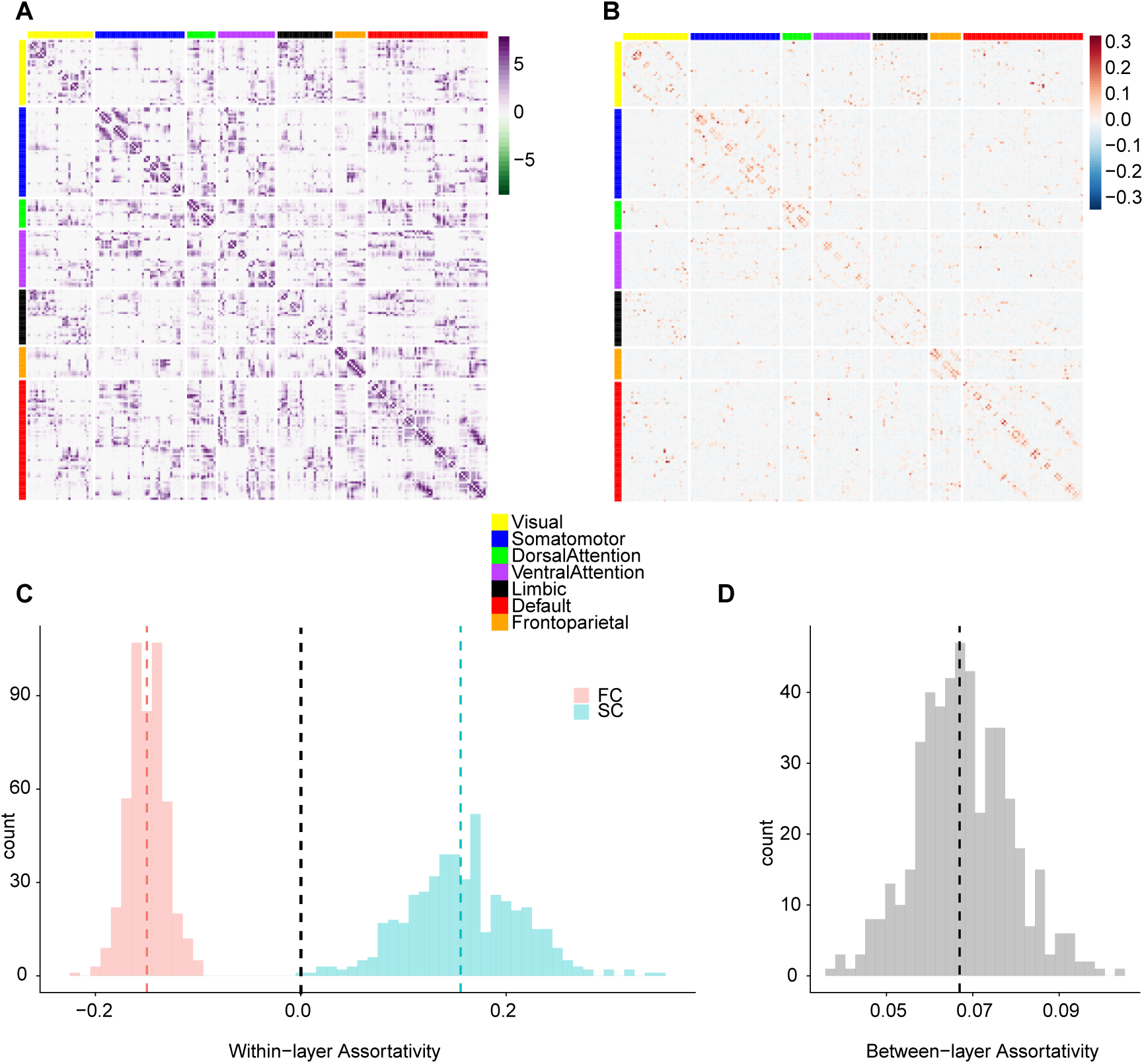
(A) Averaged SC across all subjects, with edges representing log transformed streamline counts. (B) Averaged FC across all subjects with edges representing regularized partial correlation coefficients. Nodes in both panels (A) and (B) are sorted by membership in 7 canonical resting-state networks (C) Within-layer assortativity: histograms (tallying numbers of individual subjects) of the strength assortativity within the functional network (FC) and the structural network (SC), respectively. Red: FC, Blue: SC. (D) Between-layer assortativity between FC and SC

### 3.2 Within-layer assortativity between canonical resting-state networks

Next, we investigated if these assortativity patterns were distributed homogeneously across the whole brain or whether they exhibited local order within or between subnetworks. The overall disassortative mixing in FC was more prominent within functional subnetworks derived from the canonical Yeo parcellation (Figure 3B), showing a contrast between diagonal (within a subnetwork) and off-diagonal (between subnetworks) elements. SC demonstrated overall assortative mixing between subnetworks (Figure 3A), while showing a slightly higher range of assortativity within a subnetwork (diagonal) and some more strongly disassortative mixings between certain subnetworks.

**Figure 3:**
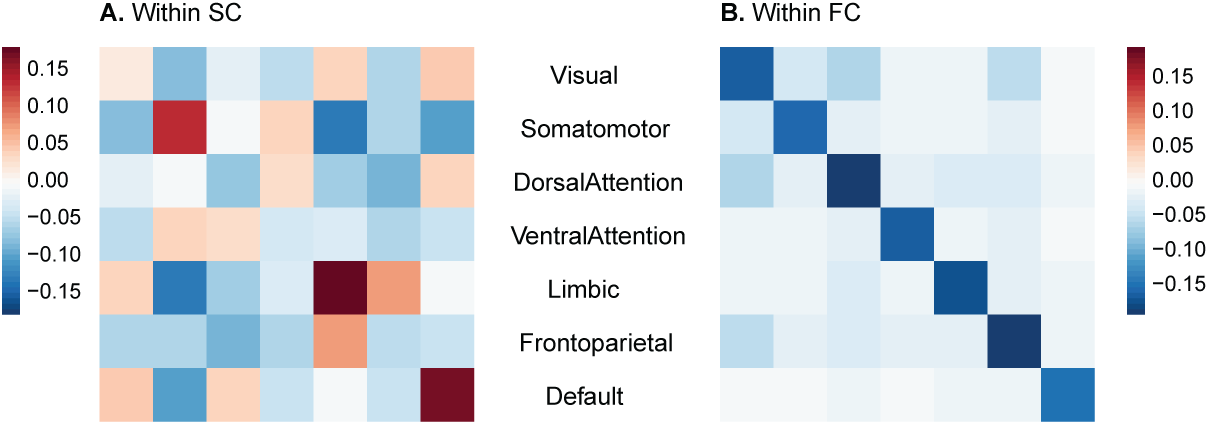
(A) Within-layer assortativity in SC between 7 canonical resting-state networks (B) Within-layer assortativity in SC between 7 canonical resting-state networks

### 3.3 Between-layer versus within-layer assortativity between 7 subnetworks

The overall assortative linkage between SC and FC could be also displayed between SC subnetworks and FC subnetworks assuming one-to-one correspondence (Figure 4). We find that the magnitude of subnetwork coupling between SC and FC ranged fairly heterogeneous across different networks.

**Figure 4:**
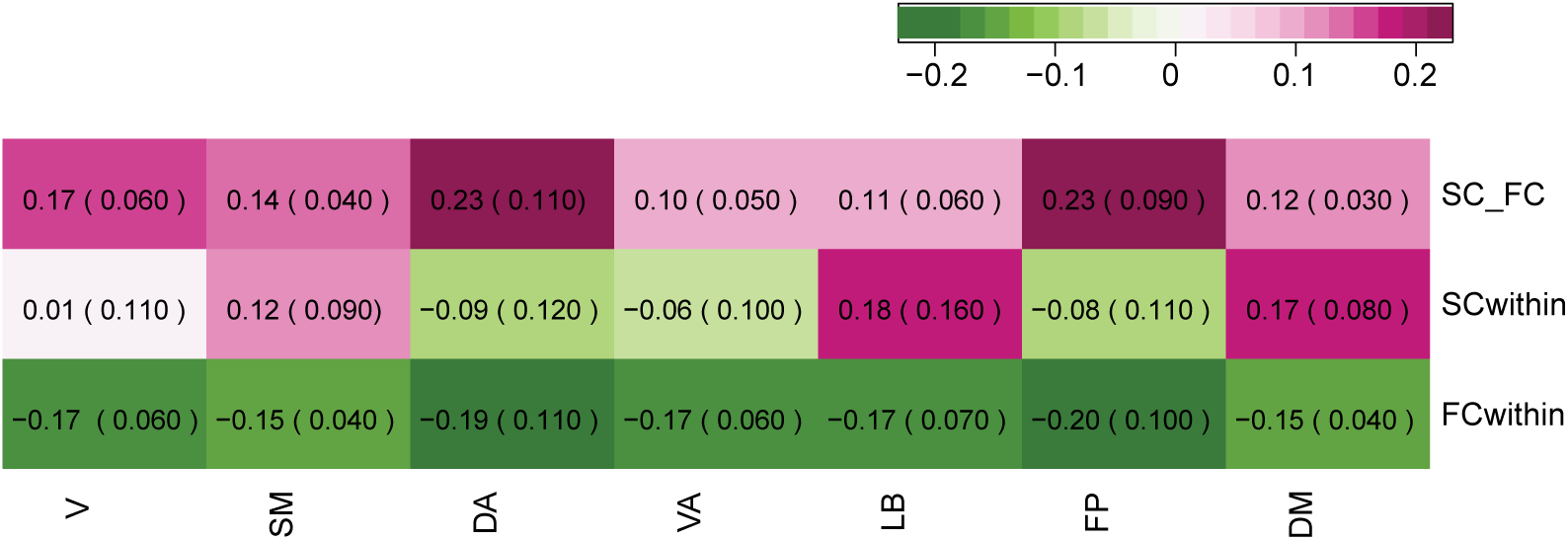
(First row) Between-layer assortativity between SC and FC, (Second row) Within-layer assortativity in SC (Third row) Within-layer assortativity in FC, numbers indicate median between-layer assortativity of the all subjects and numbers in the parentheses are median absolute deviation (MAD), V: Visual, SM: Somatomotor, DA: DorsalAttention, VA: VentralAttention, LB: Limbic, FP: FrontoParietal, and DM: Defaultmode network.

For instance, Dorsal Attention and Fronto-Parietal networks exhibited between-layer assortativity that was twice as large as compared to the rest of the subnetworks. Interestingly, those networks with higher between-layer assortativity (Dorsal Attention and Fronto-Parietal) also showed strong disassortativity within the FC layer and strong disassortativity within the SC layer. We could differentiate within-layer SC assortativity for each subnetwork, showing both assortative and disassortative mixings for subnetworks as opposed to the overall assortative mixing pattern when aggregated in a single network. Within-layer FC assortativity showed disassortativity across all subnetworks although each network exhibited a different assortativity value.

### 3.4 Left versus right hemisphere assortativity differences both within and between layers

When we further investigated the above characteristics by separating the left and the right hemispheres, we found that the characteristic disassortative mixing in within FC layer is mainly driven by the left hemisphere (Figure 5 and Figure 6). In fact, the subnetworks in the right hemisphere showed weak disassortativity within the right FC layer; Dorsal Attention and Fronto-Parietal networks still demonstrated stronger disassortativity compared to other networks but all subnetw orks in the right hemisphere showed much weaker disassortativity than those of the left hemisphere (Figure 6, all *p*-values < 10^−^21^^ after Bonferroni adjustment). In contrast, those with higher between-layer assortativity (Dorsal Attention and Fronto-Parietal) in both hemispheres displayed also strong disassortativity within the FC layer and within the SC layer in the similar way when both hemispheres were aggregated (Figure 5). We quantified the contrasts between hemispheres in both within and between-layer assortativity using permutation test based on 10000 Monte Carlo resampled approximate distribution (See methods for details). Three common features were observed for all subnetworks (Figure 6). The assortativity distributions for the left and right hemispheres showed a smaller difference between layers compared to within layer assortativity distributions except Fronto-Parietal and Default networks. Of note, the left and the right hemisphere differences showed opposite patterns between within SC and within FC; within SC, the right hemisphere was characterized with negative and smaller assortativity than those in the left hemisphere except Ventral Attention (not significant) and Fronto-Parietal networks (the trend is reversed) (Figure 6). In addition, the stark difference between hemispheres demonstrated mainly within FC. Moreover, the pattern is the opposite of within SC difference, which is summarized as box plots in the last panel of Figure 6; the left hemisphere showed strong disassortative connectivity and the right hemisphere showed weak disassortativity within the FC layer (Figure 6). However, there are interesting differences among subnetworks. For instance, unlike other subnetworks, the frontoparietal network and Default network showed small but significant strong lateralization in terms of between-layer assortativity.

**Figure 5:**
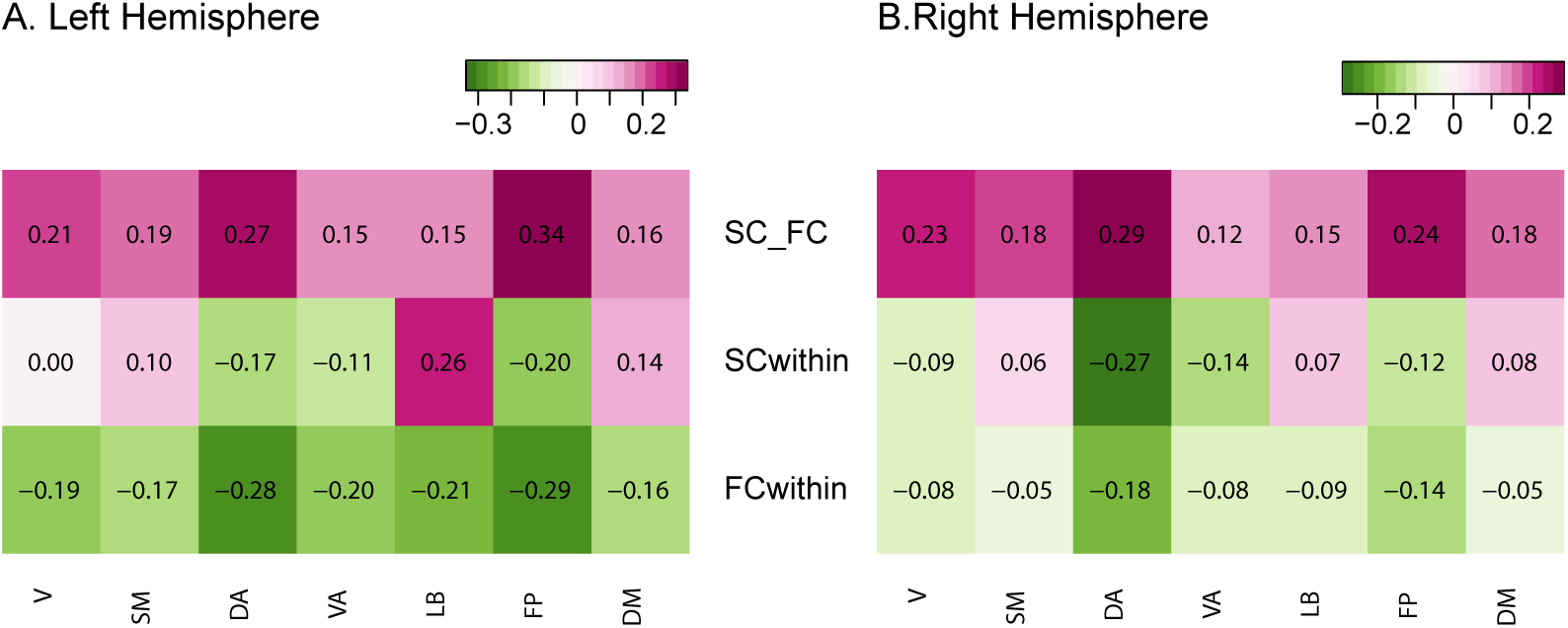
(A) Left Hemisphere, (B) Right Hemisphere. (First row) Between-layer assortativity between SC and FC, (Second row) Within-layer assortativity in SC, (Third row) Within-layer assortativity in FC, V: Visual, SM: Somatomotor, DA: Dorsal Attention, VA: Ventral Attention, LB: Limbic, FP: Fronto Parietal, and DM: Default mode network.

**Figure 6:**
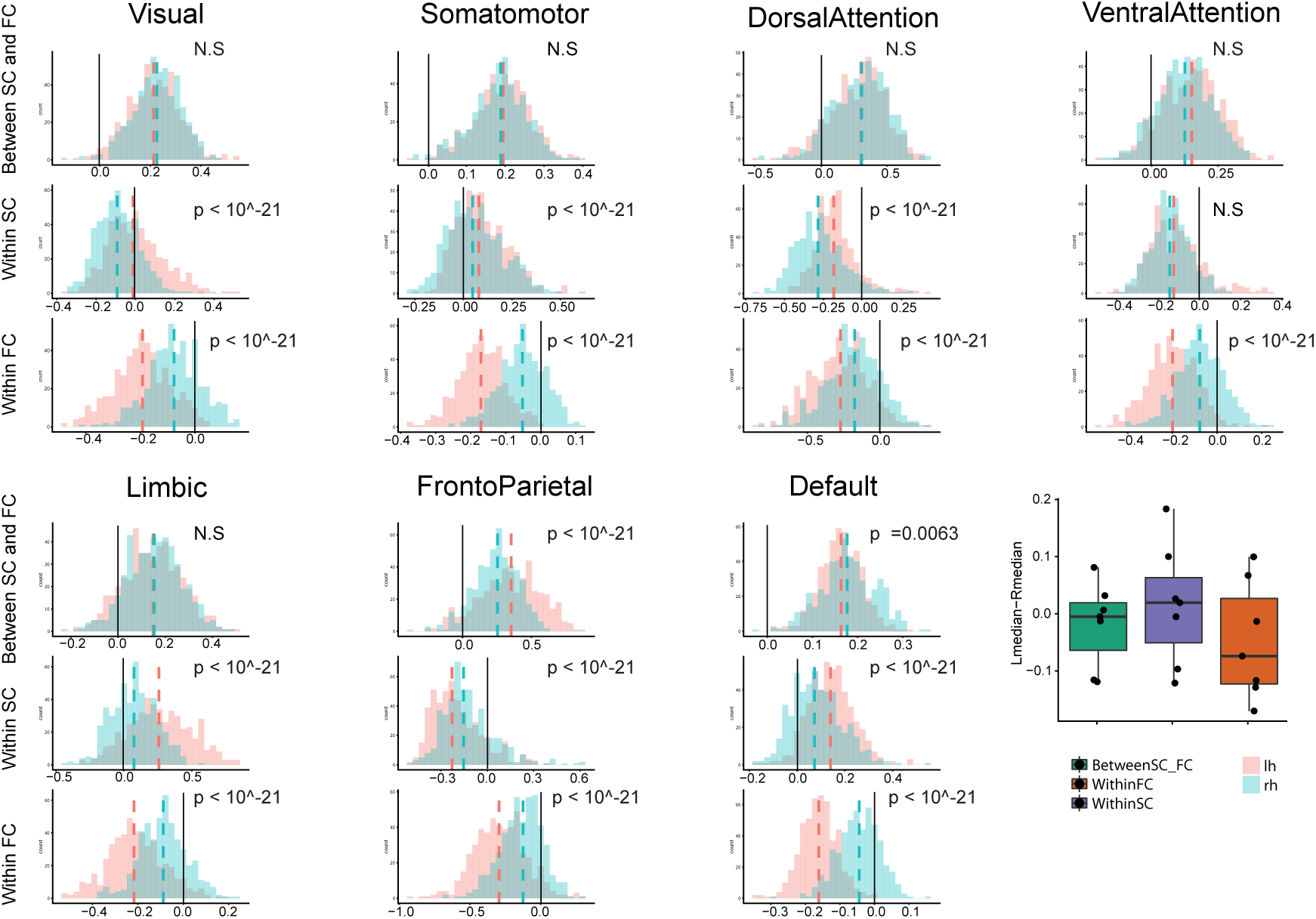
(First row) Between-layer assortativity between SC and FC, (Second row) Within-layer assortativity in SC, (Third row) Within-layer assortativity in FC (Box plots, Dashed lines indicate the medians of two distributions, Black solid line is added to identify zero on the horizontal axis. A summary box plot for the median differences between the left and the right hemispheres. X-axis: Green: between-layer assortativity, Purple: within-FC assortativity, Orange: within-SC assortativity, y-axis: the difference between the median of the left hemisphere distribution of assortativity and the median of the right hemisphere distribution of assortativity in each subnetwork.

## 4 Discussion

In this study, we constructed a two-layer multiplex interdependent network from structural and functional connectivity estimated from MRI data to examine if within- and between-layer node strength assortativity differs between SC and FC and how such differences may affect the robustness of the brain functions. In particular, we divided SC and FC into a canonical partition of seven resting state networks (RSNs) to examine within- and between-layer assortativity differences in subnetworks; furthermore, we investigated hemispheric differences to see if our findings could provide some explanations of asymmetric vulnerability and recovery process in each hemisphere. We find that, in general, SC is organized in an assortative manner, indicating brain regions are, on average, connected to other brain regions with similar node strengths. On the other hand, FC showed disassortative mixing in node strength. More detailed analysis showed that this discrepancy between SC and FC assortativity was pronounced to a different extent within- and between-RSNs. In SC, brain regions within the same subnetwork are connected with similar node strengths; in contrast, in FC brain regions are more likely connected to brain regions with similar node strengths between subnetworks rather than within its own subnetwork. In addition, these patterns showed lateralization; the overall disassortative mixing within subnetworks in FC was mainly driven from the left hemisphere. The degree of laterality also showed differences among subnetworks, which may explain why certain neurological dysfunctions mainly seem to be originated from certain subnetworks and not in the other or different recovery rate after unilateral brain injuries.

### 4.1 Assortative SC and Disassortative FC

Degree assortativity, as typically applied in network science is computed as a global network metric and typically ranges between −0.3 and 0.3 (Newman, 2003). Among biological networks, previous studies have shown disassortativity in several biological networks, including those defined by protein interactions (Newman, 2003). Instead, synaptic networks in C. elegans (Chatterjee & Sinha, 2007) and human structural brain networks estimated with diffusion imaging and tractography (Hagmann *et al*., 2008; Van Den Heuvel & Sporns, 2011)are assortative, and this assortativity appears associated with the existence of modules (Avalos-Gaytán *et al*., 2012). Functional connectivity has been reported to show assortative mixing (Eguiluz *et al*., 2005) when edges are computed as standard Pearson correlations. Assortativity in FC networks rises in the course of epileptic seizures (Bialonski & Lehnertz, 2013). To our knowledge, no previous study has examined assortative coupling within a two-layer multiplex SC/FC model. Assortative mixing in networks is known to confer greater robustness against random removal of nodes or edges compared to disassortative networks (Newman, 2003; Pechenick *et al*., 2012; Vázquez & Moreno, 2003). On the other hand, when it comes to spreading infectious diseases or seizure activity, assortativity makes it easier for these disruptions to spread across the whole network (Newman, 2003; Pechenick *et al*., 2012; Vázquez & Moreno, 2003). Against this background of theoretical work, the observed assortative and disassortative organization of SC and FC, respectively, may be important to keep brain networks intact or resilient against potential failures. For SC, to counter the effects of lesions from injuries or disease processes, having a connected network may be a priority to keep the flow of neural signals and processing as intact as possible, even if signaling paths or delays may increase due to the lesions. Hence, SC may need to be organized with positive assortativity to promote resilience. On the other hand, limiting the extent of shared neuronal information (as expressed in the statistical construct of partial correlations) or controlling the spread of abnormal brain activity such as seizures could be a significant aspect in the architecture of FC, with disassortativity helping FC to maintain its functionality against indiscriminate propagation of perturbations.

### 4.2 Between-layer assortativity between SC and FC

When more than two networks are coupled, or more generally, in a multi-layer network, the coupling between nodes in different layers affects the robustness of the system (Danziger *et al*., 2016). When nodes in different layers are connected regardless of their degrees, cascading failures of the nodes can destroy the network easily because even if a low-degree node is removed, that node can be connected to a high-degree node in another layer, and its removal could thus fragment the network into disconnected parts (Danziger *et al*., 2016; Reis *et al*., 2014). In general, when there is a positive correlation of the degree-degree coupling between layers, the interdependent networks are known to be more robust (Reis *et al*., 2014). In this study, we find that overall SC and FC are coupled in a way that nodes with similar strengths are connected between layers in all subnetworks (Figure 3 and Figure 4). This topological feature of the brain’s multi-layer organization may explain why the brain’s overall resilience against loss of nodes and edges. Theoretical studies suggest (Reis *et al*., 2014) that positive inter-layer assortativity promotes the retention of functionality unless a significant volume of the brain has been affected for example in the course of progressive neurodegenerative disease (Valenzuela & Sachdev, 2006; Yoo *et al*., 2015).

### 4.3 Subnetwork differences of SC and FC in within- and between-layer node strength assortativity

After demonstrating these topological patterns within whole-brain networks, we carried out a more detailed analysis of specific RSNs, or subnetworks, to discern if these effects were predominantly found in specific subdivisions of the cerebral cortex. We found that Visual, Somatomotor, Limbic and Default Mode Networks displayed assortativity patterns within and between layers that are consistent with robust organization. In contrast, the Dorsal Attention, Ventral Attention and Fronto Parietal networks showed disassortativity within both SC and FC layers, suggesting greater vulnerability to loss of structural network components which limits the robustness of the multi-layer network In line with previous studies, we also find distinctive differences between Dorsal and Ventral Attention networks, for instance, the between-layer assortativity of the Dorsal Attention network averaged over all participants was approximately twice as higher than that of the Ventral Attention network (Figure 5); SC within-layer assortativity in the right and the left hemispheres showed significant differences in DA but not in VA (Figure 6) (Fox *et al*., 2006; He *et al*., 2007; Vossel *et al*., 2014).

### 4.4 Lateralization of assortativity and its implication

In some cases, most notably in unilateral neglect (Heilman *et al*., 1984; Hillis, 2006; Karnath *et al*., 2004), lesions have differential effects on behavior and cognition depending on the laterality of the lesion site. Hence, we were interested to determine of our model predicted hemispheric differences in robustness. Indeed, the strong disassortativity in the FC layer observed in the whole brain mainly seems to derive from the left hemisphere, although both hemispheres show disassortativity in general within the FC layer. Stronger disassortativity in the left hemisphere suggests greater robustness to the disruptive effect of the brain injuries – conversely, weaker disassortativity in the right hemisphere suggests greater vulnerability. Increased robustness in the left hemisphere may also be followed by faster recovery post-injury. Indeed, previous studies showed ankle muscle recovery differences between the hemispheres (Kim *et al*., 2006). This pronounced lateralization of the FC layer can also be related to previous studies that showed disrupted laterality in the functional brain network in brain disorders (Swanson *et al*., 2011; Royer *et al*., 2015; Ocklenburg *et al*., 2015). The Ventral Attention network (as opposed to Dorsal Attention network) showed a large difference in the disassortativity between the left and the right hemispheres; the right hemisphere VA exhibited much weaker disassortativity than that of the left hemisphere, which is consistent with the prevalence of spatial neglect when patients experienced strokes in the right hemisphere in regions associated with the VA (He *et al*., 2007; Corbetta *et al*., 2005). We note that our findings suggest hemispheric differences in robustness despite largely symmetric distributions of standard topological measures related to both SC and FC.

### 4.5 Limitations

There are several limitations of our study. First, there are many ways to estimate functional brain networks. Depending which preprocessing steps one chooses to use, the relationship of subnetworks can vary; for instance, negative correlations between some RSNs are observed only when global signal regression is applied (Murphy & Fox, 2017). In our study, we adopted an approach to estimate partial correlations in order to allow the assessment of assortativity within a sparse functional network composed of functional links that express specific shared pairwise dependencies. More commonly used full correlation methods yield full networks and are prone to transitivity and spurious dependencies that artifactually boost shared variance. Second, assortativity could in principle be calculated based on other nodal attributes such as node between-ness or page-rank, and node strength can be also defined in different ways depending on how we define weights in FC. Third, as the assortativity is a global measure, estimating it within and between subnetworks might suffer as there are smaller numbers of nodes within each subnetwork than in the network as a whole. Future studies could investigate more detailed analyses using various nodal attributes using alternative definitions of weighted assortativity and with different parameters during time series processing.

## 5 Conclusion

In this study, we have systematically examined topological discrepancies between SC and FC by estimating node strength assortativity, using a framework of two-layer multiplex interdependent networks. We find that SC is, in general, organized as an assortative network while FC is organized as a disassortative network, with assortative coupling between the layers, an arrangement that promotes robustness within the interdependent network considered as a multi-layer system. Moreover, we find differences in subnetworks for within and between layer assortativity. Finally, we find there is a characteristic lateralization of assortativity expressed in the FC layer. Our study may be useful for quantifying the robustness of human brain networks, and for predicting individual differences in the response to injury, recovery rate or prognosis.

